# Divergence times in demosponges (Porifera): first insights from new mitogenomes and the inclusion of fossils in a birth-death clock model

**DOI:** 10.1101/159806

**Authors:** Astrid Schuster, Sergio Vargas, Ingrid S. Knapp, Shirley A. Pomponi, Robert J. Toonen, Dirk Erpenbeck, Gert Wöerheide

## Abstract

Approximately 80% of all recent sponge species belong to the class Demospongiae. Yet, despite their diversity and importance, accurate divergence times are still unknown for most demosponge clades. The estimation of demosponge divergence time is key to answering fundamental questions like e.g. the origin of Demospongiae, their diversification and historical biogeography. Molecular sequence data alone is not informative on an absolute time scale, and therefore needs to be “calibrated” with additional data such as fossils. Here, we apply the fossilized birth-death model (FBD), which has the advantage, compared to strict node dating with the oldest fossil occurrences, that it allows for the inclusion of young and old fossils in the analysis of divergence time. We use desma-bearing sponges, a diverse group of demosponges that form rigid skeletons and have a rich and continuous fossil record dating back to the Cambrian (∼500 Ma), aiming to date the demosponge radiation and constrain the timing of key evolutionary events, like the transition from marine to freshwater habitats. To do so, we assembled mitochondrial genomes of six desma-bearing demosponges from size-selected reduced-representation genomic libraries and apply a fossilized birth-death model including 30 fossils and 33 complete demosponge mitochondrial genomes to infer a dated phylogeny of Demospongiae. Our study supports a Neoproterozoic origin of Demospongiae. Novel age estimates for the split of freshwater and marine sponges dating back to the Carboniferous and the previously assumed Recent (∼18 Ma) diversification of freshwater sponges is supported. Moreover, we provide detailed age estimates for a possible diversification of Tetractinellidae (∼315 Ma), the Astrophorina (∼240 Ma), the Spirophorina (∼120 Ma) and the family Corallistidae (∼188 Ma) all of which are considered as key groups for dating the Demospongiae, due to their extraordinary rich and continuous fossil history.

## 1 Introduction

The sequencing of sponge mitochondrial (mt) genomes greatly increased in the last decade [1–5]. Nevertheless, we are still far from a representative number of mitochondrial genomes suitable as a basis for molecular phylogenetic analyses at the level of orders and below, because some key taxa, such as Demospongiae, are so far undersampled. While in the species-poorest class, Homoscleromorpha (106 species) the mt genomes for 14.2% of the species (15) are sequenced, this ratio is only 0.1-0.5% for Hexactinellida (679 species, 3 mt genomes), Calcarea (690 species, 1 mt genome), and Demospongiae (8225 species, 38 mt genomes) (Organelle Genome Resource database in GenBank; https://www.ncbi.nlm.nih.gov/genomes/OrganelleResource.cgi?taxid=6040). Therefore, there is a considerable need for denser taxonomic sequencing of mt genomes in sponges to allow for finer-scaled phylogenomic analyses.

Despite a few exceptions like *Poecillastra laminaris* (Tetractinellida: Astrophorina), where the mt genome was assembled using 454 pyrosequencing data [6], or the freshwater sponges *Spongilla lacustris* and *Ephydatia* cf. *muelleri,* which were assembled from Illumina (TruSeq) synthetic long-reads [7], all sponge mt genomes sequenced to date were assembled from sanger sequencing reads [8, 9]. However, Sanger sequencing is outdated regarding costs and yield, in particular if multiple mt genomes are pursued. Additionally, the use of this method can be challenging in (demo)sponges due to the presence of extra protein-coding genes, long intergenic regions that may include repetitive sequences [4, 10], introns in the *cox1* gene [11, 12] and the existence of different gene arrangements [3]. An extreme example of the special characteristics of sponge mitochondrial genomes is the mt genome of *Clathrina clathrus* (Calcarea, Calcinea) which encodes 37 genes distributed in six linear chromosomes ranging 7.6-9.4 kb in size [13]. Despite their somewhat unique features, mt genomes have been successfully used to infer robust demosponge phylogenies [3, 8, 9] and it is clear that gathering more sponge mt genomes will improve our understanding of the evolution of this animal group.

The demosponge order Tetractinellida comprises 23 families of world-wide distribution, of which eleven possess a rock-like skeleton built of interlocking spicules called desma. In contrast to most other demosponges, which fossil remains are usually limited to loose spicules [e.g. 14], most tetractinellid families are known for their well preserved fossils and their continuous record [e.g. 15]. Among these are the Corallistidae of which characteristic desmas (dicranoclones) are known at least since the Late Jurassic with a continuous fossil record throughout the Mesozoic and Cenozoic [15]. However, among all tetractinellids, only three complete mt genomes (i.e. *Poecillastra laminaris* [6], *Geodia neptuni* [8] and *Cinachyrella kuekenthali* [9] have been sequenced to date, none of which are from families of desma-bearing tetractinellids.

Sphaerocladina is another order of desma-bearing demosponges with a fossil record dating back to the Cambrian [15] from which no mt genome has been sequenced to date. However, this group is of particular importance for understanding demosponge evolution as it is regarded as the sister group to freshwater sponges [16–20], thus constitutes a key order for reconstructing the last common ancestor of freshwater and marine sponges.

Given the rich fossil record of these rock-sponges [21–23], sequencing the mt genomes of representatives of tetractinellids and Sphaerocladina will allow us to combine the robustness of the phylogenies inferred from mt genomes, with the rich fossil record of these sponge groups, to provide a dated phylogeny of demosponges that can be used to better understand their evolutionary history.

Here, we generated size-selected reduced representation genomic libraries [24] to *de novo* sequence and assemble the mitochondrial genomes of six species of the orders Tetractinellida (mainly Corallistidae) and Sphaerocladina. Structural features of the six novel mt genomes are discussed. In total 35 demosponge mt genomes and 30 fossil taxa of diverse ages are used to infer a dated phylogeny of Demospongiae using the fossilized birth-death (FBD) clock model. In contrast to node calibrated molecular clock models, which only allow users to set the ’oldest’ known fossil ages as constraints on certain nodes, the FBD model allows assignment of fossils of different ages to a clade without requiring morphological information about the fossils in the analysis [25]. Thus, the FBD model appears suitable for groups consisting of a rich and well studied fossil record such as desma-bearing demosponges [e.g. 21–23]. Until now the FBD model, in particular in the absence of a fossil character matrix, was used to estimate divergence times in bears [25], ferns [26], tetraodontiform fishes [27] and certain beeches [28], groups with fossils extending back to the Mesozoic. However, no attempt has been made to use this method to estimate the divergence time of groups, such as sponges, that radiated in the Early Paleozoic.

Our approach not only allows us to revise the results of previous molecular dating studies of Porifera, but also provides answers to the following questions: Can we confirm a Neoproterozoic origin of Demospongiae by our approach? When was the split of marine (Sphaerocladina) and freshwater sponges (Spongillida)? Can the suggested recent divergence time of freshwater sponges (7-10 Ma) [29, 30] be confirmed by our analysis? When was the origin of Tetractinellida and is this in congruence with the first occurrence of tetraxial-like fossil spicules?

Are the estimated divergences recovered for the suborders Spirophorina, Astrophorina and the family Corallistidae in accordance with their putative fossil appearance dates?

## 2 Materials and Methods

### 2.1 DNA extraction and Illumina library preparation

Genomic DNA was extracted using a standard phenol-chloroform protocol [31] from frozen (-80°C) sponge tissue of five species (*Corallistes* sp., *Corallistes* sp., *Neophrissospongia* sp., *Craniella* sp., *Vetulina* sp.), subsampled from the Harbor Branch Oceanographic Institute (HBOI; USA, Florida) collection and one specimen of *Cinachyrella alloclada* collected fresh and preserved at -80°C. Detailed information on the samples used including museum vouchers, location, collection date and depths is provided in the Supplementary Table 1. DNA was purified with AmpureXP (Agentcourt) beads 3-5 times according to the manufacturer protocol, to remove degenerated DNA fragments and secondary metabolites. Validation and quantification was performed using the AccuClear Ultra High Sensitivity dsDNA quantitation assay on a SpectraMax M2 plate reader (Molecular Devices, Sunnyvale, California). Two enzymes, *MboI* and *Sau3AI* (New England BioLab) were used to digest 1.0-1.3 μg DNA for 6 h at 37°C [24]. Digested products were cleaned with Ampure XP beads and eluted in 25 μl HPLC water. The Illumina KAPA Hyper Prep Kit v1.14 (Wilmington, MA) was used according to the manufacturer’s protocol for library preparation including size selection at 350-750 bp and library amplification [32]. Upon quality control (Bioanalyzer and quantitative real-time RT-PCR), all six libraries were 300 bp pair-end sequenced on an Illumina MiSeq (Illumina, Inc.) at the Hawai’i Institute of Marine Biology (HIMB) Genetics Core facility (Hawaii, USA).

### 2.2 Mitochondrial genome assembly

Forward and reverse pair-end sequences (∼2 Mio reads per library) were merged using the Paired-End reAd mergeR (PEAR) [33] software as implemented in our own Galaxy platform. A minimum overlap of 10 bp, a possible minimum length of the assembled sequences of 50 bp and a quality score threshold for the trimming of low quality parts (including adaptors and barcodes) of 20 was used. Paired sequences were imported in Geneious^®^ v8.1.8 (http://www.geneious.com, [34]) and a custom BLAST database for each library was build. For each library sequenced, one closely related mitogenome was downloaded from NCBI and used as reference genome to map mt reads against and assemble the mt genomes. The reference genome used for each species is given in Supplementary Table 1. The entire custom database was blasted against all reference genome protein, rRNA, and tRNA genes, as well as intergenic regions. To check for possible contamination, all reads were assembled separately and blasted against the NCBI database: non-sponge fragments, if any, were then excluded from the analysis. The remaining sponge sequences were mapped again to the reference genome. Possible intronic regions within the *cox1* of *Cinachyrella alloclada* were checked by blasting the library database against the *cox1* + intron region of *Cinachyrella alloclada* (HM032738). Consensus sequences were assembled *de novo* and mapped against the reference genomes respectively. Mitochondrial genomes were annotated using the similarity annotation tool (>75%) and the ORF finder as implemented in Geneious^®^ v8.1.8.

### 2.3 Protein alignment and phylogenetic reconstruction

A concatenated alignment was built using Geneious^®^ v8.1.8 from the 14 protein coding genes extracted from the mt genomes of 35 demosponge taxa. The final protein alignment was 3994 characters long, of which 1429 characters were constant, 285 characters were parsimony uninformative and 2280 characters were parsimony informative. This alignment was used to infer a Bayesian phylogenetic tree with PhyloBayes-MPI (v1.7) [35]. Two concurrent chains ran until convergence, assessed using the *tracecomp* and *bpcomp* statistics in phylobayes, with the site-heterogeneous CAT-GTR model [36]. Burn-in was conservatively set to 30% of points sampled. Additionally, a Maximum Likelihood (ML) analysis with 1,000 bootstrap replicates was done using RAxML v8.0.26 [37] and the best-fitting evolutionary model (VT+Gamma+I+F) as suggested by ProtTest 3.4 [38]; the proportion of invariant sites parameter (I) was excluded as recommended from the RAxML manual [37]. A summary tree is provided in the Supplementary Figure 1.

### 2.4 Fossils and their assignments

The protein alignment was complemented by fossil taxa and their ages (nexus file available at LMU Open Data XXX). As the FBD model requires the specification of point fossil ages [25], the youngest stratigraphic age for each fossil was taken (see Supplementary Table 2). In order to review the possible influence on the node ages of using different parameters in BEAST, we carried out two different analyses, which differ by the following parameters: 1) number of fossils, to test for the sensitivity of fossil sampling density; 2) the origin of the FBD model and the root age and; 3) the included/excluded Paleozoic fossils with sphaeroclone desmas because the homology of those spicules to the Mesozoic forms [see e.g. 39] is debatable, and to assess the impact of removing the oldest fossil on the predicted ages (see also Table 1). Fossils of 22 (BEAST analysis 1) and 30 (BEAST analysis 2) taxa belonging to five different demosponge orders (Poecilosclerida, Tethyida, Spongillida, Sphaerocladina and Tetractinellida) were extracted from the literature and linked to extant species or clades based on their suggested affinities to modern taxa (Supplementary Table 2). These also include the oldest reliable fossils known to date from Poecilosclerida (*Ophiodesia* sp., 162 Ma [40]), freshwater sponges (*Spongillina* indet., 298 Ma [41]), Sphaerocladina (*Amplaspongia bulba,* 456 Ma [42], or *Mastosia wetzleri,* 155.5 Ma [43]) and Astrophorina (*Dicranoclonella schmidti,* 150.8 Ma [44]). Detailed information for all fossils used, such as museum numbers, locality, stratigraphic level, taxonomic/systematic affinity to modern taxa, age range, references as well as the Paleobiology Database (https://paleobiodb.org/) reference number are provided in the Supplementary Table 2. Fossil taxa can be seen as either ancestor or as extinct sister taxa. Because a representative demosponge morphological data matrix is difficult to compile due to e.g. the lack of microscleres in nearly all fossils, we placed them to the appropriate subclades in the ML and BI trees (see Supplementary Figure 1). Consequently, 10 defined higher taxa of both extant and fossil sponges were constrained to be monophyletic for the analysis, namely: Tetractinellida, Sphaerocladina, Poecilosclerida, Tethyida, Haplosclerida, Spongillida, Astrophorina, Spirophorina, Corallistidae and the yet unnamed clade combining Sphaerocladina and freshwater sponges.

**Table 1:** Divergence time estimates (Ma) of demosponge clades of interest from two different anlyses. Estimates are given for the mean, and in brackets for the 95% highest posterior density interval.

| **BEAST 2.4.3 parameters** |  |  |
| --- | --- | --- |
|  | Suppl. Fig. 4<br>BEAST analysis 1<br>originFBD: **1000.0**<br>oldest Sphaerocladina fossil: **456.0**<br>number of total fossil: **22**<br>Suppl. Table 2, black colored taxa | Fig. 1<br>BEAST analysis 2<br>originFBD: **900.0**<br>oldest Sphaerocladina fossil: **155.5**<br>number of total fossil: **30**<br>Suppl. Table 2, black & red colored taxa |
| **Nodes** |  |  |
| **A** | 28 (7, 53) | 18 (5, 37) |
| **B** | 483 (467, 517) | 311 (298, 338) |
| **C** | 166 (52, 304) | 120 (40, 222) |
| **D** | 391 (232, 564) | 315 (216, 423) |
| **E** | 185 (163, 246) | 188 (155, 239) |
| **F** | 279 (178, 396) | 240 (173, 317) |
| **R** | 875 (606, 1200) | 594 (515, 730) |

### 2.5 FBD model settings

The fossilized birth-death model [25, 45] as implemented in BEAST v.2.4.3 [46] was used with the following settings for both analyses: An uncorrelated relaxed molecular clock model was chosen using default settings. No partitioning was applied on the data matrix as it had no influence on the divergence time estimation in [47] (see Table 4 and Figure 3 in [47]). For the molecular sequence data, a Gamma Site model with the JTT amino acid substitution model [48] was specified. As the start of the FBD process (root of the tree), based on previous molecular clock analyses [49, 50] we used two different ages (1000 Ma and 900 Ma) with a lognormal prior (mean=517 Ma, standard deviation min=471 Ma, max=624 Ma). Two hyperparameters were induced for the uncorrelated lognormal distribution (*ucldMean.c* and *ucldStdev.c).* As the substitution rates in Heteroscleromorpha mt genomes are considered to be low [3], we assumed an exponential prior distribution with 95% probability density on values <1 for the *ucldStdev.c* parameter. The diversification rate prior was set to an exponential with mean equal to 1.0 as the proportion of extant (33 species) and fossil taxa (22 or 30) used can be regarded as balanced. A beta distribution was chosen for the sampling proportion with Alpha 2.0. The default prior ’uniform’ (0,1) was used for the turnover parameter. Two independent Markov chain analyses were run for 400 million generations, sampling every 5000 generation. Runs were evaluated using Tracer v.1.6 [51] to assure stationarity of each Markov chain, an effective sample size (EES) for all parameters over 200, and convergence of the independent runs. The first 25% of the sampled tree topologies from both analysis were discarded as burn-in, and the remaining trees were combined in LogCombiner and summarized in TreeAnnotator (both programs are implemented in the BEAST package) with mean divergence times and 95% highest posterior density (HPD). Before this, all fossils were removed from the tree using the FullToExtantTreeConverter tool (a tool implemented in BEAUti v.2.4.3). Possible prior influences to the posterior distribution estimates were checked by specifying the sampling from the prior only and rerunning the analysis. A summarized comparison of the turnover, diversification and sampling proportion of both runs and the priors is provided in Supplementary Figure 2A-C, indicating that the number of fossils used are sufficient for our analyses. Additionally, node ages of interest from both BEAST analyses (split of freshwater sponges and Sphaerocladina, Tetractinellida, Astrophorina, Spirophorina and Corallistidae) were extracted from the combined log-output-files and histograms showing the frequency distribution of the posterior age estimates were plotted in RStudio [52, 53] (script available at openData LMU XX), indicating the 95% highest posterior density interval (HPD), the mean and the standard deviations.

## 3 Results and Discussion

### 3.1 Mitochondrial genome organisation – a general comparision

While this approach has proven useful in other taxa such as molluscs and cnidarians [54, 55], here we provide the first complete mitochondrial genomes obtained from size-selected reduced representation genomic libraries of sponges. For all six libraries, we obtained more than 2 Mio. reads of a minimum length of 50 bp and a quality score >20 for all reads. All mitochondrial genomes were circular and vary in length and GC-content between 17,364 and 20,261 bp and 32.8% to 35.7% respectively (Supplementary Figure 3), which is in line with mitogenomes of other Heteroscleromorpha [see e.g. 3]. All mitogenomes contain 24 tRNAs, 14 protein-coding genes and two ribosomal RNAs and have the same gene order and coding strand as found in their reference genomes. The mitochondrial genome of *Cinachyrella alloclada* (GW3895) contains a 1,141 bp long group I intron in the *cox1* gene, which encodes for a homing endonuclease gene (HEG) of the LAGLIDADG family (Supplementary Figure 3A). This intron is inserted at nucleotide position 723 with respect to the *cox1* sequence of *Amphimedon queenslandica* as previously found in several other species of the genus *Cinachyrella* [e.g. 11, 12]. In *Corallistes* spp. and *Neophrissospongia* sp., four gene pairs overlapped (*atp8/atp6* (1bp), *nad4L/cox1* (13bp), *nad4/trnH*(_gug_) (21bp) and *nad6/trnA_ugc_* (10bp) as previously reported for *Geodia neptuni* [8]. A further gene-pair overlap of 23 bp (*nad5/trnA(_ucg_))* was located in *Vetulina* sp. (Sphaerocladina), the same as found in freshwater sponges (e.g. *Eunapius subterraneus* and *Ephydatia muelleri)* [56]. Compared to the closest reference genome available to date (*E. subterraneus;* 88.5% pairwise sequence identity), *Vetulina* sp. is 4,589 bp shorter (total 20,261 bp) in size, shows reduced intergenic regions, lacks one tRNA gene (trnR(_ucg_)) and has two types of *trnaI* genes (trnI(_cau_) and trnI(_cau_)). The gene order and coding strands in *Vetulina* sp. is the same as for *E. subterraneus*. Although all freshwater sponges are known to possess various repeat motifs (direct, inverted and palindromes) in their mt genomes, some of which form repetitive hairpin structures [4], none of these features were found in the mt genome of *Vetulina* sp. The same applies to other assembled mitogenomes despite of their presence in other heteroscleromorphs (e.g. *Suberites domuncula* or *Axinella corrugata,* see [10]), which suggests that such repeat motifs evolved several times independently in sponges with large intergenic regions.

### 3.2 Phylogenetic analyses

Our ML and BI trees corroborate the sister group relationship of the marine order Sphaerocladina (*Vetulina*), which is morphologically characterized by the possession of sphaeroclone desmas, to freshwater sponges (Spongillida) (Supplementary Figure 1), hence strengthens previous ribosomal and partial mitochondrial single gene data [16–20]. Of the five Tetractinellida sequenced in this study, *Corallistes* spp. and *Neophrissospongia* sp. form a robust clade within the suborder Astrophorina. Furthermore, *Cinachyrella alloclada* is sister to *C. kuekenthali* and with *Craniella* sp. forming a robust clade within the subclass Spirophorina (Supplementary Figure 1). This study increases the number of currently sequenced mt genomes within the order Tetractinellida by five and supports previous phylogenies of this order based on single genes [see e.g. 19, 20, 57–59].

### 3.3 Implications for divergence time estimates for Heteroscleromorpha

The present study provides the first dated phylogeny of Heteroscleromorpha based on the relaxed molecular clock FBD model. The two analyses performed indicate a Neoproterozoic divergence of Heteroscleromorpha/Keratosa (node R, see Table 1, Figure 1, Suppl. Figure 4). As the origin time of the FBD process should be greater than the maximum value of the root age with a log-normal distribution [25], we obtained different divergence times for Heteroscleromorpha/Keratosa in both analyses with the root age affecting the divergence time (see Table 1, Figure 1, node R). Previous ages estimated for crown-group Demospongiae, using different software, clock-model settings, and taxon sets, varied between 657-872 Ma [30, 47, 50, 60], which is in the range of both of our analyses (see Table 1, Figure 1 and Suppl. Figure 4, node R). The first reliable fossil representing crown-group Demospongiae was described by [61] from the early Cambrian (515 Ma). As this fossil only constitutes a minimum age it does not contradict a possible Neoproterozoic divergence of demosponges. This deep origin of crown-group Demospongiae concurs with the first appearance of demosponge-specific biomarkers (24-ipc sterol) in rocks dating 540-650 Ma (Neoproterozoic) and today present in all major demosponge clades [49, 62]. Additional paleontological evidence for an early divergence of sponges is given by the discovery of a 600 Ma old fossil, with poriferan features and interpreted as a stem group descendant for all sponges [63].

**Figure 1:**
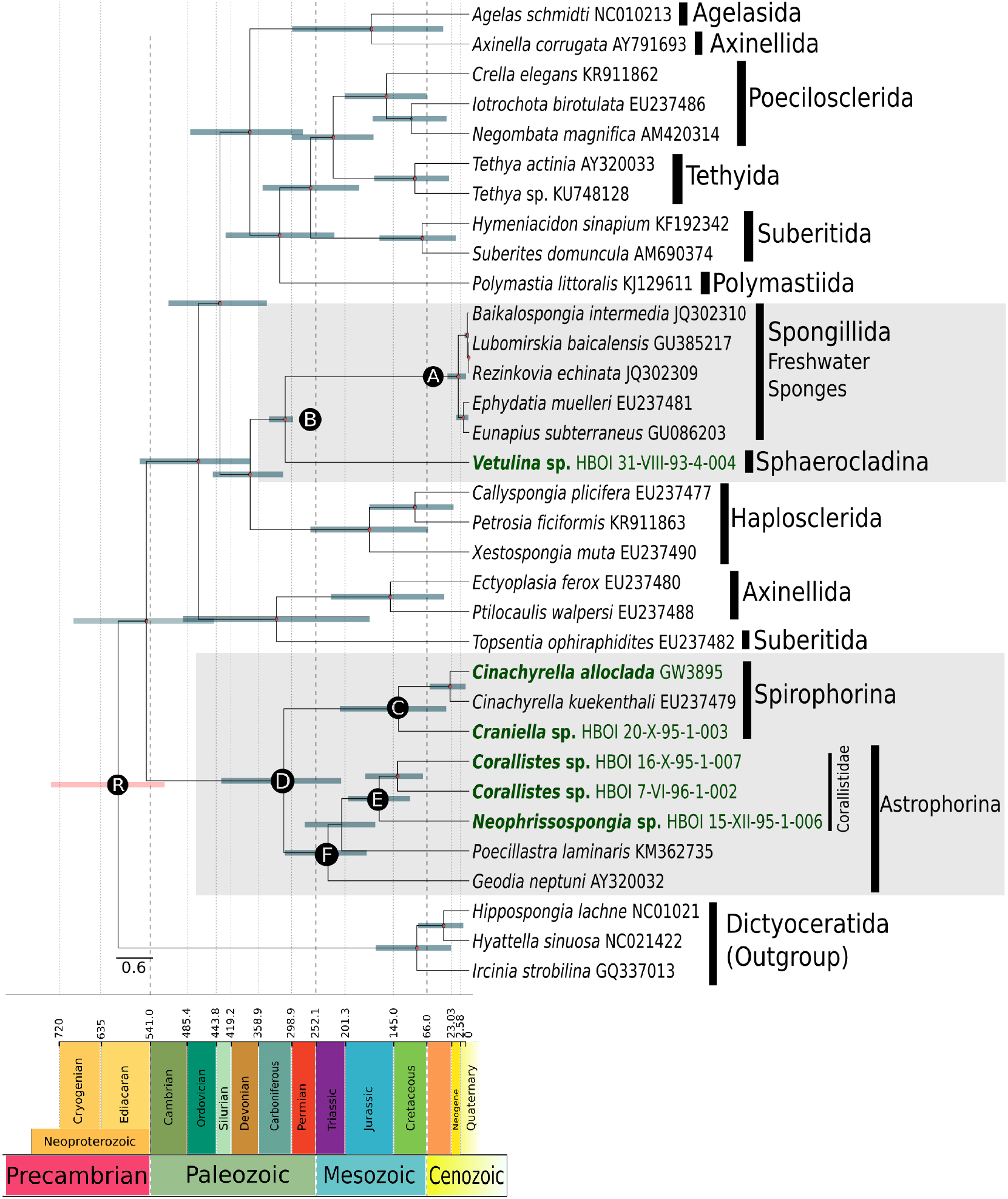
Time calibrated phylogeny of Demospongiae. Based on paramters of BEAST analysis 2 plotted on stratigraphic chart. New sequenced species are in dark green and bold. Taxonomic clades of interest are shaded in light gray. Error bras on node ages are in dark turquoise. Nodes of interest are marked with a capital letters A-F and correspond to node ages listed in Table 1. The capital letter R specifies the root age of the dated phylogeny.

### 3.4 Inferred divergence scenarios for the split of marine and freshwater sponges

Many earlier molecular dating studies of Porifera were based on mitogenomic datasets, however, these were hampered by an incomplete taxon sampling, for example lacking freshwater and desma-bearing sponges. Consequently, inferences of divergence times for key demosponge taxa, such as the split between marine and freshwater sponges could not be addressed. Now, with the complete mitogenome of *Vetulina* sp. (Sphaerocladina) available, this study represents the first dated phylogeny that suggests a likely timeframe for the split of marine and freshwater sponges. Hypothesizing that the oldest fossil with sphaeroclone desmas from the Paleozoic (*Amplaspongia bulba,* Upper Ordovician ∼456 Ma) [42] resembles species with the same desma types as those found in the Mesozoic, although larger in sizes and more heavily silicified [e.g. 64], our analysis dates the split between marine and freshwater sponges (Point B, Suppl. Figure 4) to the Early Ordovician (∼483 Ma). Sponges with a massively silicified sphaeroclone desma skeleton are well known in the Paleozoic, and were common during the late Ordovician, Middle Silurian and late Devonian [see e.g. 39, 64–67]. But, no sphaeroclone desmas are reported from the Carboniferous until the Middle Jurassic, which represents a ∼200 Ma gap in the fossil record [e.g. 64]. Due to this long gap, it is debatable whether the Paleozoic sphaeroclone desmas are homologous to those found in the Mesozoic and Cenozoic [64, 68], and therefore suitable as fossil constraint. If Paleozoic sphaeroclone desmas are excluded from the analysis (BEAST analysis 2), the mean age of the split between marine and freshwater sponges dates back to the Carboniferous and is ∼172 Ma younger (see node B in Figure 1, Table 1). The lack of a fossil sphaeroclone desmas during the Carboniferous until the Middle Jurassic has been proposed to be connected to the Permian-Triassic boundary (PTB) mass extinction [39], which led to the reduction in size of sponge spicules, the disappearance of certain sponges groups [69] and to the habitat displacement of several sponge taxa from shallow neritic environments to deeper bathyal waters [see e.g. 70]. [Maldonado et al. 71] proposed that the observed decline and turnover of the sponge fauna in the Mesozoic resulted from the reduction of silica in the oceans. This hypothesis is corroborated by the lack of sphaeroclone desmas found around and past the PTB mass extinction as well as the observed change from massive-large sphaeroclones in the Paleozoic to smaller and less silicified sphaeroclones in the Mesozoic.

### 3.5 Inferred timing of extant freshwater sponge diversification

The occurrence of the earliest freshwater sponge fossil spicule is dated to the Permo-Carboniferous [41] and constitute the first and only known fossil record of freshwater sponges from the Paleozoic. The radiation of recent freshwater sponges, however, is dated as much younger in both of our analyses (18.0-28.3 Ma, Paleogene, Table 1, Node A). Therefore, our results question [41]’s interpretation as Paleozoic spicules. Also, [72] interpreted the findings of [41] as either marine or marine influenced, which again challenge the interpretation of this oldest described freshwater sponge. In contrary, fossil freshwater sponges with intact gemmules (i.e. freshwater-sponge specific buds for asexual reproduction highly resistant to desiccation, freezing and anoxia [73, 74]) are well-known from the lower Cretaceous [75], thus supporting a diversification of Recent freshwater sponges before the Paleogene (66 Ma). Yet, [30] suggested a divergence of 7-10 Ma for Recent freshwater sponges using a node-calibrated relaxed molecular clock approach, whereupon the study of [76] indicate a Paleogene divergence.

The Paleogene record of freshwater sponges is known to be more diverse than the Neogene record [77, 78]. Our analysis includes three freshwater species (*Baikalospongia intermedia, Lubomirskia baicalensis* and *Rezinkovia echinata,* all Lubomirskiidae), all of which are known to be endemic to Lake Baikal [29, 79]. Our dated phylogeny suggests a divergence of this clade to the Early Pliocene (∼3.4 Ma, Figure 1, Node A), which correlate to the known fossil record from this area (3.2-2.8 Ma) [29, 79]. As gemmules are known from the fossil record since the lower Cretaceous [75], and are present in the Recent spongillids *Ephydatia* and *Eunapius,* but absent from Lubomirskiidae [80], our data is consistent with the hypothesis that the most recent common ancestor of Spongillida possessed gemmules, which were subsequently lost in several endemic lineages such as the Lake Baikal Lubomirskiidae [see discussion in 80].

### 3.6 Inferred divergence scenario of Tetractinellida, Spirophorina and Corallistidae

We estimated a mean origin age for Tetractinellida of 315 Ma (Late Carboniferous) (node D, Table 1, Figure 1, BEAST analysis 2), with a normal frequency distribution on the node age (Figure 1, BEAST analysis 2). Indeed, a Carboniferous origin is late for this group considering previous estimates which point to a Middle Cambrian (∼514 Ma) origin of this clade in addition to the earliest tetraxial-like fossil spicules known from the Middle Cambrian (510-520 Ma) [47, 81]. Despite these Cambrian fossil discoveries, the molecular clock analyses of [50] (∼385 Ma) and [76] (∼345.7 Ma) provide support for a post-Cambrian origin of this clade. These contradictory results may have different reasons. First, due to their massive and thicker sizes the Cambrian tetractinellid (tetraxial-like) spicules may not be homologous to post-Paleozoic forms [e.g. 39, 64]. Second, the presence of aster-like and monaxon spicules in several recent demosponge groups other than the Tetractinellida may lead to the erroneous interpretation of the Cambrian fossil spicules. Third, the high level of secondary losses of various spicule types in particular of microscleres within Astrophorina [20, 57, 82] hamper unambiguous interpretation of their homology.

The astrophorid family Corallistidae (node E, Table 1, Figure 1), characterized by dicranoclone desmas, is here dated to ~188.7 Ma (Lower Jurassic). The node age shows a left-skewed distribution to younger ages (Suppl. Figure 5, BEAST analysis 2), which correlates with the current known fossil record from the late Jurassic to Recent [15, 23]. Additional support for a Jurassic origin of the included Recent tetractinellids is provided by a node-based calibrated single-gene phylogeny (*cox1*) of [76], who dated Corallistidae to ~155 Ma. The only known fossil representative of the genus *Neophrissospongia* is described from the Early Campanian of Poland [23], but our analysis indicates a deeper origin dating back to the Middle Jurassic (Figure 1). As this family shows one of the richest and continuous fossil records among the included taxa, we tested this clade for sampling sensitivity of the FBD clock model by increasing the number of fossils by 50% (Suppl. Table 2, BEAST analysis 2). This increase of the fossil sampling neither influenced our results positively (by reducing the error bars for instance) nor negatively, which corroborates other findings of [25] and [26]. The investigation of the divergence ages of this desma-bearing demosponge family strengthen the Jurassic origin of this clade and provides additional information on possible calibration constraints for further molecular clock approaches.

The tetractinellid suborder Spirophorina (node C, Table 1, Figure 1, BEAST analysis 2) is dated to ∼120 Ma (Late Cretaceous). The frequency distribution on the node age indicates a slightly right-shifted normal distribution (Suppl. Figure 5, BEAST analysis 2). A characteristic diagnostic feature for this group is the presence of so called sigmaspire (S-to C-shaped) microclere spicules [83]. [84] described a C-shaped microsclere [Plate 24 in 84] from the Middle Cambrian Daly and Georgina Basin (Northern Territory in Australia, see Plate 24 in [84]), which he associated to “orthocladine” sponges. [85] suggested the occurrence of Spirophorina in the Early Paleozoic, with a possible Cambrian origin, however, these observations can not be supported by any of our analyses. As sigma-like spicules are also present in other demosponge lineages like e.g. in Poecilosclerida and Desmacellida, the discovered C-shaped microsclere described in [84] might not be homologous to those of Spirophorina.

## 4 Conclusion

We here successfully assembled six complete mitogenomes of different demo-sponge taxa generated by a size-selected reduced representation genomic library. Integrating these data into a novel mitogenome alignment in tandem with a newly tested relaxed molecular clock approach based on the FBD model, we provide new insights into the evolution of selected Demospongiae. The Neoproterozoic origin of Demospongiae is confirmed. Furthermore, the origin and diversification of the Tetractinellida is dated to ∼315 Ma, the suborders Astrophorina to ∼240 Ma, the Spirophorina to ∼120 Ma and the family Corallistidae to ∼188 Ma. Furthermore, we discovered that increasing the fossil sampling by 50% within the Corallistidae indicates that this approach is relatively insensitive to fossil sampling density, which corroborates with the findings of other studies [25, 26]. Nevertheless, our estimated divergence times of different higher tetractinellid taxa such as the Astrophorina or Corallistidae can be further used for inferring finer-scaled divergence time estimates to shed new light on e.g the correlations of secondarily spicule losses to possible changing geochemical/geological historical events in the past.

The split of freshwater sponges and marine Sphaerocladina is dated to ∼311 Ma, most of which correlate with the fossil record. Additionally, we confirmed previously assumed recent (∼18 Ma) diversification of freshwater sponges. These results, and in particular the dated split of freshwater and marine sponges, can be used as a root age for further dated phylogenies on freshwater sponges in order to get a better idea of e.g. their historical biogeographical processes such as the radiation timing in different ancient lakes.

## Acknowledgements

Funding was provided by the German Research Foundation (DFG) (grant number DFG ER 611/3-1, DFG Wo896/15-1) and the LMU Mentoring Program. Samples were collected by HBOI’s Johnson-Sea-Link manned submersible during several epeditions. We greatly acknowledge Zac H. Forsman for discussions on library preparation and Amy Eggers for sequencing (Hawaii Institute of Marine Biology). We are thankful to Amy Wright for sharing samples and Megan Conkling for helping in the collection (Harbor Branch Florida Atlantic University, HBOI).

